# A deep-learning-based toolbox for Automated Limb Motion Analysis (ALMA) in murine models of neurological disorders

**DOI:** 10.1101/2021.05.27.445999

**Authors:** Almir Aljovic, Shuqing Zhao, Maryam Chahin, Clara de la Rosa del Val, Valerie Van Steenbergen, Martin Kerschensteiner, Florence M Bareyre

**Author notes:** Contributed equally (Almir Aljovic, Shuqing Zhao). **Correspondence should be addressed to:** Florence M. Bareyre.

## Abstract

In neuroscience research, the refined analysis of rodent locomotion is complex and cumbersome, and access to the technique is limited because of the necessity for expensive equipment. In this study, we implemented a new deep-learning-based toolbox for Automated Limb Motion Analysis (ALMA) that requires only basic behavioral equipment and an inexpensive camera. The ALMA toolbox enables the unbiased and comprehensive analyses of locomotor kinematics and paw placement and can be applied to neurological conditions affecting the brain and spinal cord. We demonstrated that the ALMA toolbox can (1) robustly track the evolution of locomotor deficits after spinal cord injury, (2) sensitively detect locomotor abnormalities after traumatic brain injury, and (3) correctly predict disease onset in a multiple sclerosis model. We, therefore, established a broadly applicable automated and standardized approach that requires minimal financial and time commitments to facilitate the comprehensive analysis of locomotion in rodent disease models.

## Introduction

Research into neurological conditions often attempts to uncover how the structural and functional deficits of individual neurons and circuits relate to behavioral outcomes^1^. Over the years, a number of elaborate tools based on chemogenetics, optogenics, and connectomics have been designed to manipulate and record the structure and function of individual circuits and circuit elements^2–6^; nevertheless, a full understanding of the consequences of such interventions can only be achieved using refined behavioral analysis.

To this extent, a wide range of behavioral tests have been developed that can detect specific aspects of behavior in a range of neurological conditions^7–9^. Although such tests have substantially improved our ability to selectively monitor deficits, for example in motor function, the evaluation tools are still hampered by a number of limitations. These include a lack of automatization, which makes such tests both highly time-consuming and susceptible to observer bias; the high level of investment needed for the recording and analysis equipment, which often restricts access to the technique to a few specialized labs; and the limited scope and sensitivity of the analyses, as they are often based on a set of pre-selected parameters. Such limitations also affect the evaluation of locomotion that plays a central role in disease-related neuroscientific research, as many common neurological conditions caused by trauma, ischemia, or inflammation prominently affect walking abilities.

Unraveling the complexity of locomotion is best approached via the generation of gait parameters based on the precise position of the limbs. Since Hildebrand’s early description of the notion of gait^10^, several tests have been designed to evaluate limb motion and paw placement. Early work relied on collecting footprints^11–13^ from an animal after inking the paws, but the data that could be retrieved was often incomplete due to rapid drying of the ink, and analysis was cumbersome to perform. Later systems, including the use of the commercial automated Catwalk® system, which uses foot placement to derive a range of gait parameters that reflect locomotion on a static ramp, offered important improvements^14,15^. The Catwalk® system has proven quite useful in the analysis of gait parameters in rodents subjected to genetic manipulation or injuries^16,17^. However, this system also has some limitations, as the tracking of light-furred rodents can be difficult, and potential mistakes in the tracking of footprints need to be manually corrected by the experimenter. In addition, the system is static and does not allow the locomotion speed to be controlled. The gait can also be recorded at variable speeds while the animal is running on a treadmill using three-dimensional video recordings coupled with a kinematic tracing system^13^ or using the well-established motion-capture system VICON with eight infrared cameras^18,19,20^. Both require reflective markers to be manually and bilaterally attached at key joints (for example, iliac crest, lateral malleolus, the tip of the toe, etc.), and use of the latter system is further limited by the cost of the acquisition^21^. Therefore, recent new advances have been made to improve and facilitate locomotion tracking. A key step has been the development of the DeepLabCut (DLC) method that provides a markerless approach to labelling and tracking moving joints (DeepLabCut^22^) and, thereby, facilitates the kinematic analysis of gait as well as arm movements in animals and humans^23–26^. However, even this approach requires substantial specialized expertise and processing times to translate the limb coordinates into kinematic profiles and parameters that quantify the distinct aspects of locomotion.

Here, we describe how we overcame this final challenge by developing an open-source computational “toolbox”, Automated Limb Motion Analysis (ALMA), which provides fully automated and comprehensive analysis of locomotion and fine paw placement in mice with minimal costs, time requirements, and previous expertise. This toolbox includes a graphical user interface (GUI) with functionalities for automated kinematic parameter computation and automated footfall detection, kinematic data analysis with random forest classification and principal component analysis, and visualization of the gait kinematics. To make it amenable to as many users as possible, we used only inexpensive equipment (custom-made ladders and a commercial treadmill) and a single standard high-speed action camera. We applied this toolbox to mouse models of common traumatic and inflammatory diseases of the brain and spinal cord to demonstrate its capability to robustly and sensitively monitor the evolution of locomotor deficits in a broad range of neurological conditions. Notably, our results show that such an automated comprehensive analysis can delineate the specific parameters of locomotor function that are best suited to track injuries of the brain or spinal cord or are sensitive enough to predict disease onset during the prodromal phase of a multiple sclerosis model.

## Results

### Pipeline for automated limb motion analysis: DLC markerless labeling, model training, and automated kinematic analysis and footfall detection in mice

In this paper, we present a new open-source toolbox for the analysis of gait kinematic parameters of locomotion and fine paw placement in mice. To do so, we focused on two behavioral tasks that were applied to mice: walking on a treadmill (**Fig. 1a**) to determine gait kinematic parameters and walking along ladders (**Fig. 1b**) with regularly or irregularly spaced rungs to analyze fine paw placement. All tasks were filmed with a GoPro 8 camera positioned parallel to and at a fixed distance and angle from the treadmill and ladder. We used DLC to perform markerless limb tracking of six hindlimb joints (toe, metatarsophalangeal joint [MTP], ankle, knee, hip, and iliac crest) for treadmill kinematic analysis, and of limb extremities for the ladder rung paradigm. (**Fig. 1a,b**). For both analyses, we focused only on the hindlimb trajectories.

**Figure 1:**
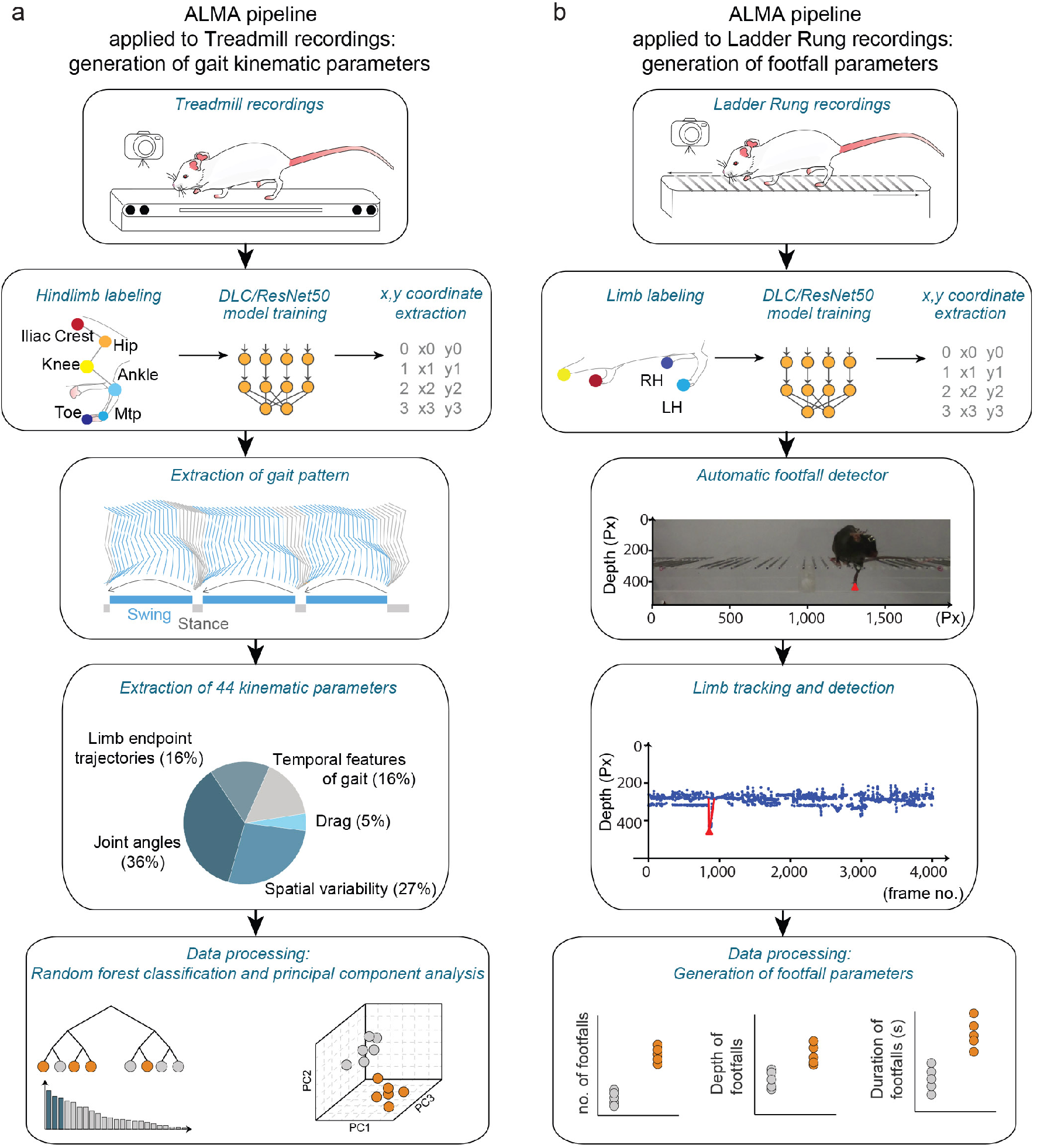
Automated behavioral analysis using ALMA toolbox. **a**- Schematic of ALMA toolbox application to treadmill recordings for the generation of gait kinematic parameters. First, treadmill videos were recorded with a GoPro 8 camera placed parallel to the treadmill. Then, markerless limb labeling and modeling was trained and refined using ResNet-50 and DeepLabCut (DLC) and coordinates were extracted. Coordinates were tracked for six hindlimb joints (toe, metatarsophalangeal [MTP] joint, ankle, knee, hip, and iliac crest) in the treadmill kinematic paradigm, and the coordinates were processed using the toolbox to generate hindlimb trajectories. This allowed the generation of 44 kinematic parameters that represented joint angles, spatial variability, limb endpoint trajectories, temporal features of gait, and dragging. Data processing was conducted in ALMA to obtain principal component analysis and random forest classification of the parameters. **b**- Schematic of ALMA toolbox application to the ladder task recordings for the generation of footfall parameters. First, ladder task videos were recorded with a GoPro 8 camera placed parallel to the ladder. Then, markerless limb labeling and modeling was trained and refined using ResNet-50 and DLC, and coordinates were extracted. Coordinates could be tracked for all four paws. The toolbox used the automated footfall detector to extract limb tracing and footfall detection. The on- and off-set of locomotor errors (footfalls) were estimated using signal processing methods and subjected to manual validation. Three parameters, the number, depth, and duration of the footfalls, were extracted. Px: Pixels

We then trained the neural network model for body part coordinate extraction as described in the methods. For each behavioral paradigm, 2D body part coordinates were obtained using a deep residual network model (ResNet-50), which was trained and refined using DLC (**Fig. 1a,b**). We then performed automated kinematic analysis on the coordinate data obtained from the treadmill task and extracted 44 kinematic parameters that reflected the locomotion of the hindlimbs, which were grouped into five main categories: joint angles, spatial variability, temporal feature of gait, limb endpoint trajectories, and dragging (**Fig. 1a and Extended Table 1**). These parameters could then be classified and compared using random forest and principal component analyses (**Fig. 1a**). For the ladder rung analysis, we performed automated detection of hindlimb footfalls (**Fig. 1b**), which were manually validated. Three parameters were then extracted, the number, depth, and duration of footfalls (**Fig. 1b**).

### Tracking locomotion after spinal cord injury with the ALMA toolbox

We first tested the applicability of our toolbox for analyzing overground locomotion following spinal cord injury (SCI) (**Fig. 2a**). To do so, mice walking on a treadmill were recorded with a GoPro 8 camera (**Fig. 2b**), and we used DLC to perform markerless labeling of the six hindlimb joints as described above (**Fig. 2c and Extended Video 1**). The ALMA toolbox then automatically detected the treadmill speed and adjusted the body part coordinates accordingly. The adjusted body part coordinate data were then used to extract stride onsets and offsets, as well as to distinguish the stance and swing phases (**Fig. 2d and Extended Fig. 1**). This procedure enabled automated kinematic parameter extraction for each stride (**Fig. 2d**). We validated the reliability of the extracted data with baseline recordings to check the reproducibility of the 44 parameters measured. To do so, we tested and re-tested mice on consecutive days on the treadmill task at baseline (e.g., before injury), and we found that reliability was excellent for all parameters measured (**Fig. 2e**; *r* = 0.9985, *p* < 0.0001, Pearson’s correlation) between the initial test and the re-test conditions. For the SCI experiment we used mice that were first tested in the Basso Mouse Scale (BMS)^27^ and showed at least one point of recovery over time (BMS score baseline 9±0; BMS score 3dpi 3.86±1.08 and BMS score 21dpi 6.29±0,81). We then analyzed the treadmill recordings collected before spinal cord injury and at 3 and 21 days following SCI in these mice and generated the adjusted hindlimb trajectories (**Fig. 2f**). Those trajectories were profoundly altered in SCI mice at 3 days post-injury (dpi) but had partially recovered by 21 dpi (**Fig. 2f**). We applied random forest classification to the data for the animals based on the 44 extracted parameters, and we could predict the injury status to a 98% accuracy when we compared baseline to animals at 3 dpi, and the recovery status to a 94% accuracy when we compared 3 dpi to 21 dpi animals. The step height, knee joint flexion, and knee joint extension showed the highest Gini impurity-based feature importance (**Fig. 2g**). We then compared the gait of the mice before and after spinal cord injury using principal component analysis of our 44 kinematic parameters and determined that the data for injured mice at 3 dpi clustered separately from those of the mice at baseline, while mice at 21 dpi were clustered in between, again indicating incomplete recovery of locomotor function (**Fig. 2h**). PC1 and PC2 together represented 58% of the variance, with PC2 better reflecting the between-group differences (**Fig. 2h**). After calculating the factor loadings, we identified three parameters that best represented PC2 and were, thus, most likely to track SCI related differences (**Fig. 2h**). Step height, knee joint extension, and dynamic time warping (DTW) demonstrated significant alterations at 3 dpi, and recovered over time until they were no longer significantly different from baseline (**Fig. 2h and Extended Table 2**). Our analysis thus suggests that these kinematic parameters are best suited to monitor locomotor deficits resulting from an incomplete spinal cord injury. Importantly those changes are specific of SCI as our toolbox could not pick up any gait kinematic changes following sham injury (**Extended Fig. 2**).

**Figure 2:**
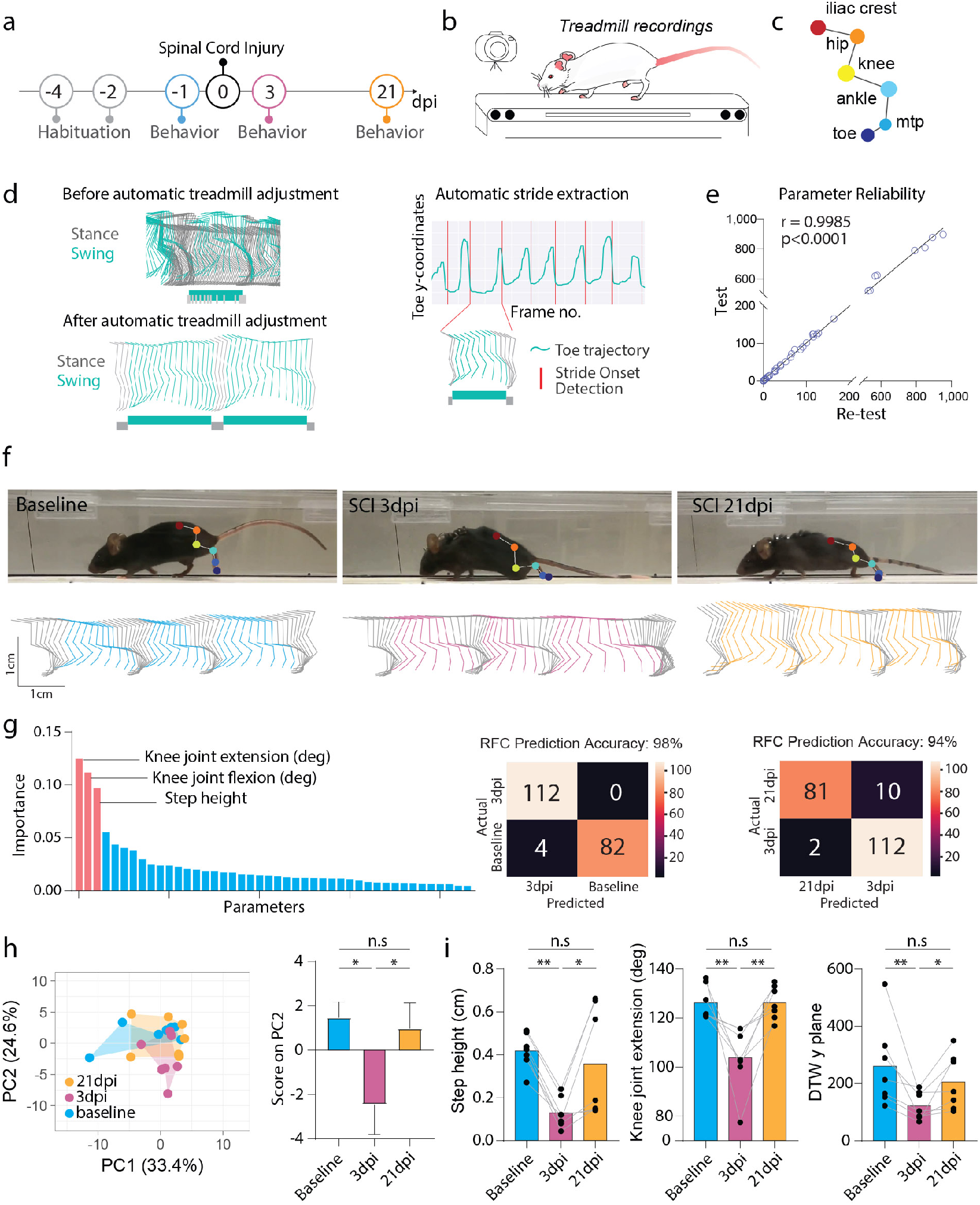
ALMA analysis of gait changes in spinal cord injured mice tested on the treadmill. **a-** Timeline of the traumatic spinal cord injury (SCI) experiment. **b-** Schematic of the treadmill system used to record the behavior of mice during the SCI experiment. **c-** Schematic of DeepLabCut (DLC) markerless joint labeling. Six joints were labeled: iliac crest, hip, knee, ankle, metatarsophalangeal joint (MTP), and toe. **d-** Representation of hindlimb trajectories (left panel) before adjustment in ALMA (top; green, swing; grey, stance) and after adjustment with ALMA (bottom; green, swing; grey, stance). Note, that after adjustment, swing and stance were efficiently separated. Representation of automatic stride extraction from the toe coordinates and frame number (right panel). **e-** Quantification of parameter reliability; baseline data were tested and re-tested and demonstrated a high correlation coefficient (r = 0.9985, p < 0.0001; Pearson’s correlation coefficient). **f-** Photographic images (top) of mice running on the treadmill showing the markerless labeling of hindlimb joints using DLC at baseline, 3 dpi, and 21 dpi, and hindlimb trajectories for baseline (cyan), 3 dpi (fuchsia), and 21 dpi (orange). **g-** Random forest classification (RFC) of 44 parameters extracted from the ALMA toolbox for the analysis of gait following spinal cord injury and accuracy injury status prediction based on the 44 parameters using confusion matrices for 3 dpi vs. 21 dpi (Gini impurity-based feature importance for RFC: knee joint extension, 0.125; knee joint flexion, 0.112; step height, 0.097. RFC prediction accuracy: baseline vs. 3 dpi 98% and 3 dpi vs. 21 dpi 94%, tested in n = 84-92 step cycles). **h-** Principal component analysis of data obtained on the treadmill and processed with the ALMA toolbox for spinal cord injury, and plot of scores of PC2 that represent 22.3% of the variability (principal component analysis, PC1 36.1%, PC2 22.3%, repeated measures one-way ANOVA followed by Tukey’s test; baseline vs. 3 dpi [p = 0.022]; baseline vs. 21 dpi [p = 0.920], 3 dpi vs. 21 dpi [p = 0.044]; n = 7). **i-** Quantitative evaluation of parameters associated with PC2, such as step height, knee joint extension, or dynamic time warping (DTW) y plane, at baseline, 3 dpi, and 21 dpi. Repeated measures one-way ANOVA followed by Tukey’s test was used to analyze knee joint extension (baseline vs. 3 dpi, p = 0.005; baseline vs. 21 dpi, p > 0.999; 3 dpi vs. 21 dpi, p = 0.005; n = 7), Friedman and Dunn tests were used for DTW y plane (baseline vs. 3 dpi, p = 0.004; baseline vs. 21 dpi, p > 0.999; 3 dpi vs. 21 dpi, p = 0.049; n = 7), Repeated measures one-way ANOVA followed by Tukey’s test was used to analyze step height (baseline vs. 3 dpi, p = 0.063; baseline vs. 21 dpi, p = 0.012; 3 dpi vs. 21 dpi, p > 0.999; n = 6). In all panels, data are presented as mean± SEM; *p < 0.05; **p < 0.01; ***p < 0.001. Px: Pixels

### Tracking skilled paw placement after spinal cord injury with the ALMA toolbox

While overground locomotion strongly depends on the function of intraspinal circuits, skilled paw placement requires supraspinal input and is, thus, commonly used to assess the regeneration and remodeling of descending motor tracks, including corticospinal projections. We, therefore, assessed whether the ALMA toolbox could also be used to monitor skilled paw placement following spinal cord injury (**Fig. 3a**). To this end, we recorded mice walking on a horizontal ladder with regularly or irregularly spaced rungs. Similar to the overground locomotion experiment, all videos were recorded using a single GoPro 8 camera, and we used DLC to perform markerless labeling of the hind paws (**Fig. 3b and Extended Video 2**). We then applied ALMA to determine footfall characteristics with a peak detection algorithm (**Fig. 3c,d and Extended Fig. 3**). The parameters extracted from ALMA showed that spinal cord injury caused an increase in the mean number and depth of footfalls at 3 dpi, while the mean duration of the footfalls was only slightly increased, for the ladder with regularly spaced rungs (**Fig. 3e**). Those parameters remained altered over a prolonged period, so that they remained different from baseline at 21 dpi (**Fig. 3e**). Similarly, in the ladder with irregular rungs experiment, the mean number of footfalls and duration of footfalls were found to increase at 3 dpi and remained elevated till 21 dpi. However, the mean number of footfalls significantly recovered from 3 dpi to 21 dpi. The mean depth of footfalls was, in this case, unchanged throughout the study period (**Fig. 3f**). We also calculated the total duration and total depth of the footfalls on the ladders with regularly and irregularly spaced rungs and observed a significant increase in the total depth and total duration at 3 dpi and 21 dpi (**Fig. 3g**). Finally, we compared the ALMA automated detection of the number of footfalls using a deviation algorithm with the manual correction provided by the GUI and with a fully manual analysis of the videos and in both cases found a highly significant correlation (**Fig. 3h**) confirming that the ALMA toolbox can provide a precise fully automated quantification of skilled paw placement.

**Figure 3:**
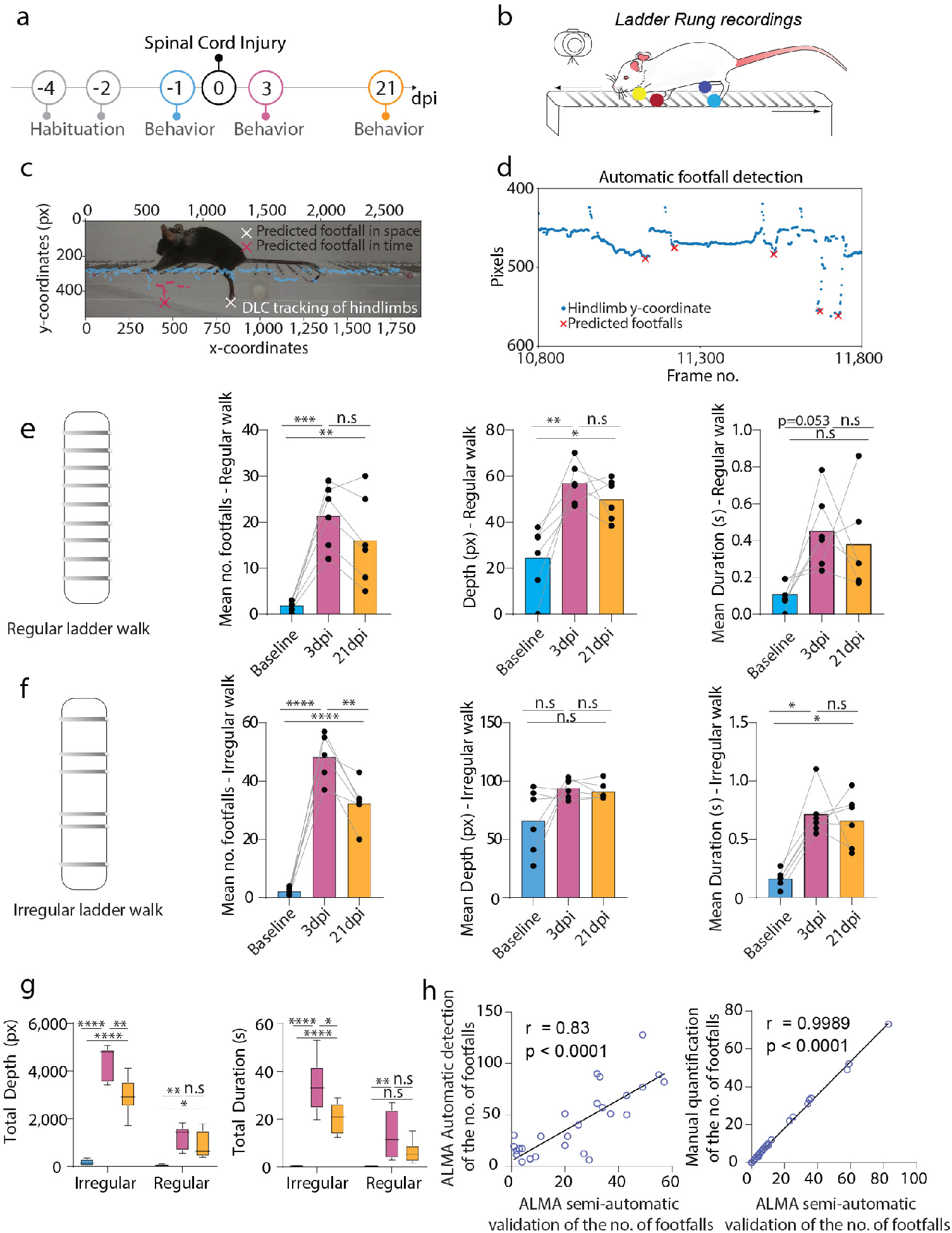
ALMA analysis of fine paw placement in spinal cord injured mice in the ladder rung test. **a-** Timeline of the traumatic spinal cord injury experiment. **b-** Schematic of the ladder rung system used to record the fine paw placement of mice during the spinal cord injury experiment indicating the DeepLabCut (DLC) markerless paw labeling (yellow, red, dark blue and light blue dots). **c-** Photographic image of a mouse running on the treadmill showing the automatic detection of footfall, as predicted in time and space by the toolbox. **d-** Automated detection algorithm used to predict footfalls in space and time. **e-** Quantitative parameters extracted from ALMA for the regular walk showing the mean number, mean depth, and mean duration of footfalls for all time points (cyan, baseline; purple, 3 dpi; and orange, 21 dpi). Repeated one-way ANOVA followed by Tukey’s test was used to the analyze the regular ladder rung results (mean no. footfalls, baseline vs. 3 dpi [p = 0.0002], baseline vs. 21 dpi [p = 0.021], 3 dpi vs. 21 dpi [p = 0.223]; mean depth, baseline vs. 3 dpi [p = 0.003], baseline vs. 21 dpi [p = 0.013], 3 dpi vs. 21 dpi [p = 0.612]; and mean duration, baseline vs. 3 dpi [p = 0.053], baseline vs. 21 dpi [p = 0.131], 3 dpi vs. 21 dpi [p = 0.838]; n = 6). **f-** Quantitative parameters extracted from ALMA for the regular walk showing the mean number, mean depth, and mean duration of footfalls for all time points (cyan, baseline; purple, 3 dpi; and orange, 21 dpi). Repeated one-way ANOVA followed by Tukey’s test was used to the analyze the irregular ladder rung results (mean no. footfalls, baseline vs. 3 dpi: [p < 0.0001], baseline vs. 21 dpi [p < 0.0001], 3 dpi vs. 21 dpi [p = 0.002]; mean depth, baseline vs. 3 dpi [p = 0.745], baseline vs. 21 dpi [p > 0.999], 3 dpi vs. 21 dpi [p >0.999]; and mean duration, baseline vs. 3 dpi, p = [0.028], baseline vs. 21 dpi [p = 0.028], 3 dpi vs. 21 dpi [p > 0.999]; n = 6). **g-** Quantitative evaluation of the total depth and total duration of footfalls for all time points (cyan, baseline; purple, 3 dpi; and orange, 21 dpi). Repeated one-way ANOVA followed by Tukey’s test was used to analyze total footfall depth on regular ladder rungs (baseline vs. 3 dpi, p = 0.0047; baseline vs. 21 dpi, p = 0.043; and 3 dpi vs. 21 dpi, p = 0.175; n=6), total depth on irregular ladder rungs (baseline vs. 3 dpi, p < 0.0001; baseline vs. 21 dpi, p < 0.0001; 3 dpi vs. 21 dpi, p = 0.0039; n=6), total duration on regular ladder rungs (baseline vs. 3 dpi, p = 0.0042; baseline vs. 21 dpi, p = 0.173; 3 dpi vs. 21 dpi, p = 0.101; n=6), and total duration on irregular ladder rungs (baseline vs. 3 dpi, p < 0.0001; baseline vs. 21 dpi, p = 0.004; 3 dpi vs. 21 dpi, p = 0.041; n = 6). **h-** Correlation between the ALMA automatic detection of the number of footfalls using the deviation algorithm and the ALMA semi-manual validation (left panel; r = 0.81, p < 0.0001; Pearson’s correlation coefficient) and between the ALMA automatic detection of the number of footfalls using the deviation algorithm and a fully manual detection (left panel; r = 0.9989, p < 0.0001; Pearson’s correlation coefficient). In all panels, data are presented as mean± SEM; *p < 0.05; **p < 0.01; ***p < 0.001. Px: Pixels; dpi: days post-injury.

### The ALMA toolbox can reveal behavioral consequences of traumatic brain injury

Next, we wanted to determine whether our toolbox, which was initially designed to reveal motor abnormalities related to spinal cord pathology, was also capable of detecting the locomotor changes induced by neurological conditions primarily affecting the brain. This is of particular importance for traumatic brain injuries, as it has often been challenging to detect and reliably quantify motor deficits arising from such insults, in particular in the acute injury phase^28,29^. We, therefore, induced moderate brain injury in mice using a traumatic brain injury (TBI) impactor and recorded the locomotion and paw placements of mice walking on the treadmill and ladders, respectively, before and 1 day and 10 days after traumatic brain injury (**Fig. 4a**). We applied the ALMA toolbox as described above and initially analyzed the treadmill recordings (**Fig. 4b,c**). The findings showed that the limb trajectories were strongly affected at 1 dpi and had partially recovered by 10 dpi (**Fig. 4d**). We applied random forest classification to all animals at baseline and 1 dpi based on the 44 extracted parameters, and we were able to demonstrate the parameters with the highest Gini impurity-based feature importance (step height, hip joint, hip joint amplitude and hip joint flexion) and predict the injury status with 83% accuracy. This indicates that our toolbox makes it possible to robustly detect even the subtle locomotor changes induced by a moderate TBI (**Fig. 4e**). We then used principal component analysis to reduce the dimensionality of our data and determine the individual movement parameters that best identified differences between the groups. We found that data for mice at baseline clustered closer to the data for the mice at 10 dpi than to those at 1 dpi, indicating that the injury and recovery effects can be ascertained based on the kinematic parameters (**Fig. 4f**). As PC1 represented almost 40% of the variance, we plotted the principal component analysis scores and demonstrated the above-mentioned effects at 1 dpi (**Fig. 4f**). Based on PC1, we identified the top three parameters that contributed to the differences: step height, stride length, and DTW distance. These parameters showed significant changes at 1 dpi and later recovery at 10 dpi, indicating that our toolbox can also detect changes in locomotion following brain lesions (**Fig. 4g and Extended Table 3**).

**Figure 4:**
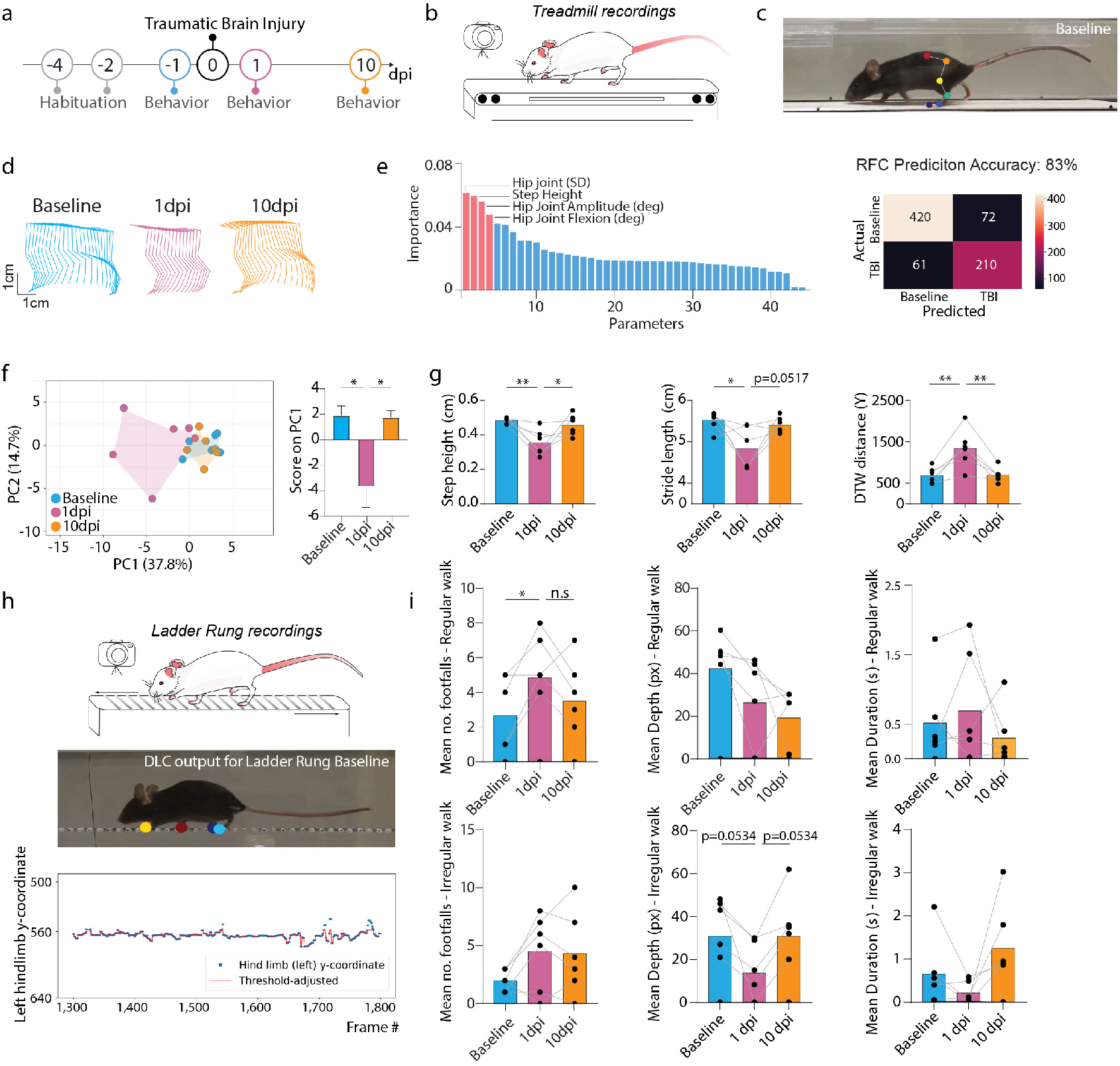
ALMA monitoring of gait changes and differences in fine paw placement in brain injured mice. **a**- Timeline of the traumatic brain injury experiment. **b**- Schematic of the treadmill system used to record the behavior of the mice during the traumatic brain injury experiment. **c-**Photographic images of the mice running on the treadmill showing markerless labeling of hindlimb joints using DeepLabCut (DLC) at baseline. **d-** Hindlimb trajectories for baseline (top, cyan), 1 dpi (middle, purple), and 10 dpi (bottom, orange). **e-** Random forest classification (left) of the 44 parameters extracted from the ALMA toolbox for the analysis of gait following traumatic brain injury, and confusion matrix (right) for determining pre-diction accuracy of the injury status based on the 44 parameters (Gini impurity-based feature importance for RFC: hip joint, 0.061; step height, 0.059; hip joint amplitude, 0.056; hip joint flexion, 0.047: RFC prediction accuracy: baseline vs. 1 dpi 83%; tested in n = 282-481 step cycles). **f-** Principal component analysis of data obtained from the treadmill task and processed with the ALMA toolbox, and plot of PC1scores that represent 37.8% of the variability and associated factor loadings (principal component analysis, PC1 37.8%, PC2 14.7%; repeated one-way ANOVA followed by Tukey’s test, baseline vs. 1dpi [p = 0.012], baseline vs. 10 dpi [p = 0.665], 1 dpi vs. 10 dpi [p = 0.014]; n = 6). **g-** Quantitative evaluation of factors associated with PC1, i.e., step height, stride length, and dynamic-time warping (DTW) distance, at baseline, 1 dpi, and 10 dpi. Repeated one-way ANOVA followed by Tukey’s test was used to analyze step height (baseline vs. 1 dpi, p = 0.0095; baseline vs. 10 dpi, p = 0.730; 1 dpi vs. 10 dpi, p = 0.033; n=6), stride length (baseline vs. 1 dpi, p = 0.0214; baseline vs. 10 dpi, p = 0.855; 1 dpi vs. 10 dpi, p = 0.0517; n=6), and DTW distance (baseline vs. 1 dpi, p = 0.001; baseline vs. 10 dpi, p = 0.9943; 1 dpi vs. 10 dpi, p = 0.0012; n = 6). **h-** Schematic of the ladder rung system used to record the behavior of mice during the traumatic brain injury experiment, and photographic images of a mouse running on the treadmill showing markerless labeling of hindlimb paws using DLC at baseline and showing the algorithm detection of footfall. **i-** Quantitative evaluation of three parameters extracted from ALMA for footfalls at baseline, 1 dpi, and 10 dpi. Friedman followed by Dunn’s test was used to analyze the regular ladder rung mean no. footfalls (base-line vs. 1 dpi, p = 0.0315, baseline vs. 10 dpi, p = 0.2557; 1 dpi vs. 10 dpi, p = 0.1583), mean depth (baseline vs. 1 dpi, p = 0.1299; baseline vs. 10 dpi, p = 0.0628; 1 dpi vs. 10 dpi, p > 9999), and mean duration (baseline vs. 1 dpi, p > 0.9999; baseline vs. 10 dpi, p > 0.9999; 1 dpi vs. 10 dpi, p > 9999; n = 6) and the irregular ladder rung mean no. footfalls (baseline vs. 1 dpi, p = 0.3371; baseline vs. 10 dpi, p = 0.9370; 1 dpi vs. 10 dpi, p > 0.9999; n=6), mean depth (baseline vs. 1 dpi, p = 0.0534, baseline vs. 10 dpi, p >0.9999; 1 dpi vs. 10 dpi, p = 0.0534; n=6) and mean duration (baseline vs. 1 dpi, p > 0.9999; baseline vs. 10 dpi, p = 0.9370; 1 dpi vs. 10 dpi, p = 0.3371; n = 6). In all panels, data are presented as mean± SEM; *p < 0.05; **p < 0.01; ***p < 0.001. Px: Pixels; dpi: days post-injury.

We then analyzed the ladder task recordings using the ALMA toolbox (**Fig. 4h**). We extracted the three footfall parameters described previously and demonstrated that mice with brain injuries displayed an increased number of footfalls in the regular-rung ladder task, which returned to baseline at 10 dpi (**Fig. 4i**). In contrast, the irregular-rung ladder performance was unaffected by moderate traumatic brain injury. Taken together, these results indicate that the ALMA toolbox can be applied to assess neurological conditions of the brain and is capable of sensitively tracking both alterations in locomotor kinematics and foot placement after TBI.

### The ALMA toolbox can monitor disease symptoms and predict disease onset in a multiple sclerosis model

Finally, we sought to investigate the applicability of our toolbox for assessing neurological conditions with a less predictable disease course, such as those caused by CNS inflammation. Therefore, we induced a commonly used mouse model of multiple sclerosis, experimental autoimmune encephalomyelitis (EAE), by immunizing mice with the myelin oligodendrocyte glycoprotein (MOG). We then recorded the locomotion of the mice on the treadmill at baseline, disease onset (defined as the first day with clinical symptoms), disease peak (3 or 4 days after onset), and recovery (10 or 11 days after onset; **Fig. 5a,b**). As expected, as the disease progressed, the mice developed ascending paresis and paralysis that strongly altered their hindlimb trajectories at the peak of the disease (and precluded the use of the ladder tests). While the hindlimb trajectories recovered slightly over the next few days, they remained clearly distinct from the baseline pattern (**Fig. 5c**). Principal component analysis confirmed that the data formed distinct clusters at the onset, peak, and recovery of disease compared to baseline (**Fig. 5d**). As PC1 represented 76.1% of the variability, we plotted the scores in PC1 to recapitulate the evolution of motor systems (**Fig.5e**) and extracted the key parameters clustering with PC1. Those parameters (stride length, stride height, toe-crest distance, and dragging percentage) all showed significant changes at the peak of the disease and tended towards a later recovery, indicating that our toolbox is well suited to monitoring the locomotor alterations resulting from the formation and resolution of inflammatory lesions (**Fig. 5f and Extended Table 4**).

**Figure 5:**
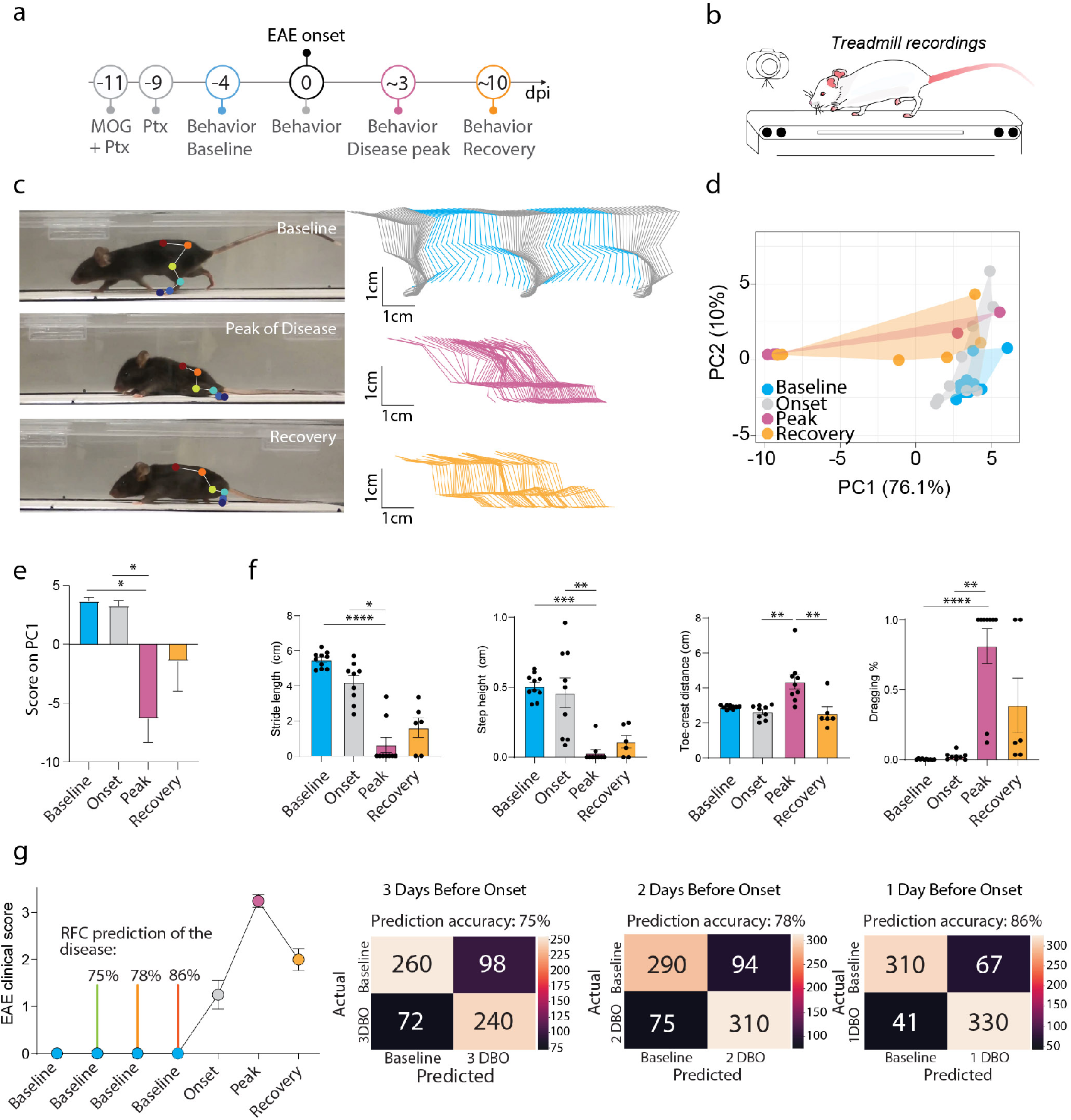
ALMA monitoring of locomotor changes in mice that developed experimental autoimmune encephalomyelitis and accurate prediction of disease development during the prodromal phase. **a**- Timeline of the EAE experiment. **b**- Schematic of treadmill system used to record the behavior of the mice during the EAE experiment. **c**- Photographic images of mice running on the treadmill (left) showing markerless labeling of hindlimb joints using DeepLabCut (DLC) at baseline (top), onset of disease (middle), and disease recovery (bottom), and hindlimb trajectories (right) for baseline (top), onset of disease (middle), and disease recovery (bottom). **d**- Principal component analysis of data obtained on the treadmill and processed with the ALMA toolbox. **e**- Plot of the PC1 scores that represent 76.1% of the variability and associated factor loadings (Kruskal–Wallis followed by Dunn’s test; baseline vs. peak, p = 0.0104; onset vs. peak, p = 0.0418; baseline vs. recovery, p= 0.9806; peak vs. recovery, p > 0.9999). **f**- Quantitative evaluation of factors associated with PC1, i.e., stride length, step height, toe-crest distance, and dragging, at baseline and different stages of EAE (Kruskal–Wallis followed by Dunn’s test; stride length, baseline vs. peak [p < 0.0001], onset vs. peak [p = 0.0231], baseline vs. recovery [p = 0.0019], peak vs. recovery [p > 0.9999]; n=6; step height, baseline vs. peak [p = 0.0002], onset vs. peak [p = 0.0020], baseline vs. recovery, [p = 0.0382], peak vs. recovery [p >0.9999]; toe-crest distance, baseline vs. peak [p = 0.0737], onset vs. peak [p = 0.0036], baseline vs. recovery [p = 0.0382], peak vs. recovery [p = 0.7374]; and dragging (%), baseline vs. peak [p < 0.0001], onset vs. peak [p = 0.0082], baseline vs. recovery [p = 0.0039], peak vs. recovery [p > 0.9999]; n=6). **g**- EAE clinical score and prediction of the disease onset based on random forest classification in the prodromal phase (3, 2, and 1 days before onset) using ALMA. In all panels, data are presented as mean± SEM; *p < 0.05; **p < 0.01; ***p < 0.001. Px: Pixels; dpi: days post-injury.

We next examined whether this refined analysis of locomotion kinematics is sufficiently sensitive to pick up the prodromal stages of the disease, which are not apparent in classical EAE scoring (based on simple observations of mouse mobility). In this context, we were particularly interested in whether we could detect the more subtle locomotor alterations associated with the initial formation of the lesions, which can predict the subsequent onset of overt disease symptoms (defined as the first day of detectable symptoms in the EAE score; often seen as tail paralysis). This would be useful, as it is often important to precisely initiate interventions, e.g., pharmacological treatment, when CNS lesions first start to form, as, once initiated, the CNS inflammatory process can self-perpetuate. Therefore, we used the ALMA toolbox to analyze the treadmill recordings starting 3 days prior to disease onset (defined by the EAE score). Using random forest classification, we found we could predict whether and when the mice would subsequently show EAE symptoms with 75% accuracy 3 days before onset, with 78% accuracy 2 days before onset, and with 86% accuracy 1 day before onset (**Fig. 5g**). This illustrated the ability of our toolbox to accurately predict whether an immunized mouse will develop overt disease symptoms in the prodromal disease phase and even predict the day of onset. This renders the initial formation phase of CNS lesions amenable to study.

## Discussion

The quantitative assessment of locomotor performance is critical in the analysis of physiological gait patterns and their perturbations in neurological diseases. Conventional evaluation strategies often rely on resource-intensive and time-demanding observation and analysis setups. Together with the specialized expertise required to operate these setups, these demands currently prevent the broad application of refined locomotor analyses in basic and clinical neuroscience. In this study, we used deep-learning strategies to implement ALMA, an open-source toolbox that facilitates refined analysis of overground locomotion and skilled paw placement at a fraction of the cost of specialized behavioral setups. ALMA is fully automated, saving time and preventing observer bias, and it can be used without previous expertise through its user-friendly interface. Therefore, ALMA makes comprehensive analysis of locomotion accessible to every research group interested in revealing the behavioral consequences of nervous system dysfunction and disease.

The use of machine learning approaches to the study of behavior in both rodents and humans has dramatically increased in the last 5 years^22–25,30^. An important step in this process was the development, by Mathis and colleagues, of a markerless method (DLC^22^) to label the joints and follow their motion reliably over time. Here, we made use of this method to mark important joints in mice and import the DLC coordinates to track locomotion and fine paw placements. Alternatively, our toolbox can also be efficiently used with coordinates generated through other techniques, such as VICON or the newly published DANNCE^26^ methods.

To translate these limb and joint coordinates into locomotor patterns on a treadmill or horizontal ladder and to subsequently extract the parameters of gait and footfall, we used model training. In healthy mice, this allowed us to detect up to 44 hindlimb gait parameters with almost perfect re-test reliability (r = 0.9985). It is remarkable that not only can this highly robust kinematic analysis be achieved with a fully automated analysis pipeline but also that it can be achieved with the use of an affordable single action camera. Particularly as, to date, scientists have had to rely on the use of high-end cameras and software modules to manually extract gait cycles and parameters in the analysis of kinematic behavior. Previous techniques came with several pitfalls, including high software costs, limited troubleshooting support, and cumbersome and time-consuming post-experimental processing. In contrast, the ALMA toolbox simply requires the import of DLC output using a GUI and allows the extraction of the final gait kinematic parameters. Another important aspect of our approach is that the ALMA toolbox allows the tracking of locomotor patterns based on side-view camera recordings, which, while harder to model, provide much more gait parameters compared with the bottom-view recordings used in the catwalk approach or in recent automated approaches^17,25^. As demonstrated here, 2D side-view tracking and analysis provides important information on each joint angle, step height, and body support, which is missing in top- or bottom-view recordings of gait analysis.

In addition to kinematic analysis, the ALMA toolbox offers the first automated analysis of the number, duration, and depth of footfalls on regularly and irregularly spaced ladder rungs. This is an important improvement for those investigating skilled paw placement, as it replaces a cumbersome analysis protocol that had to be performed frame by frame^31,32^. The analysis of skilled paw placement is complementary to the obtention of kinematic parameters, as it provides a readout of the damage and recovery of supraspinal circuits^16,33^. Finally, it should be noted that while we focused our analysis on both the gait and footfall aspects of mouse hindlimb movements, the ALMA toolbox should (with minor modifications) be equally suitable for the analysis of forelimb function, forelimb–hindlimb coordination, or the tracking of locomotor function in larger rodents such as rats.

We further showed that the ALMA toolbox can be applied to a range of neurological conditions and enables the automated tracking of locomotor deficits in mice with traumatic and inflammatory injuries of the brain and spinal cord. Using spinal cord injury models, we confirmed that tracking of locomotor deficits is robust, as demonstrated by the high reproducibility and the correct prediction of injury status. As spinal cord injuries lead to pronounced locomotor deficits, a large number of studies have used behavioral testing strategies to assess the functional outcomes of genetic or therapeutic modulations^18,19,34–40^. The application of the ALMA toolbox in such models should both improve the reliability of the analysis (as observer and selection biases are removed) and provide a more refined view of locomotor disturbance and recovery, as it allows the operator to precisely determine which component or components of locomotor function are regulated by a specific circuit modulation or therapeutic intervention.

The locomotor deficits observed after spinal cord injury are pronounced compared with the less obvious deficits resulting from mild to moderate brain injuries. It has often been challenging to detect and reliably monitor locomotor deficits following such brain injuries in mice, particularly in the early phase when the effects are subtle^28,29^. Our findings showed that the kinematic tracking of gait parameters, in particular, step height, stride length, and DTW (which measures the similarity between two step sequences), allowed us to sensitively reveal both the emergence of locomotor deficits (1 day after injury) and their recovery (by 10 days after injury). Interestingly, while pronounced abnormalities of locomotion were observed at the initial recording after injury, the animals later recovered a movement pattern almost identical to the pattern observed at baseline. This is clearly distinct from the comparably incomplete recovery process observed after spinal cord injury and may indicate that the persistence of intraspinal circuits is critical for re-establishing the “original” physiological movement pattern, while the reorganization of intraspinal (but not supraspinal) circuits results in a compensatory adaption of the movement pattern.

In comparison to studies of the injured brain and spinal cord, only a few attempts have been made to apply refined behavioral testing to models of inflammatory CNS damage, such as the EAE model^41–43^. One limitation has been, that, due to the disseminated nature of inflammatory infiltration, which results in variable neuronal tracking systems being affected in different mice, inter-individual differences in the pattern of locomotor changes are to be expected. For this purpose, targeted EAE models have been developed that allow inflammatory lesions to be directed to a predetermined anatomical location in the brain or spinal cord^44–46^. However, the stereotactic injection procedure required for the induction of such models limits their application, e.g., for studying disease initiation. Here we were able to show that the comprehensive kinematic analysis generated by the ALMA toolbox allows for the refined monitoring of locomotor deficits, even in classical disseminated EAE models. While such locomotor symptoms can also be tracked by the classical EAE scoring scale, which is based on visual inspection of the walking abilities of mice, kinematic analysis provides a number of advantages. First, the ALMA toolbox allows for fully automated and standardized analyses, removing observer bias and, thereby, making behavioral assessments more comparable between different observers and labs. Second, it stands to reason that the quantitative assessment of 44 distinct gait parameters should be more sensitive to differences in locomotor function compared with visual inspection. This is particularly important when evaluating the behavioral consequences of therapeutic manipulations targeting neuronal protection and repair^47^ that are expected to lead to more subtle changes in locomotor function. Third, and related to the latter, our current analysis showed that the ALMA toolbox is able to accurately predict the onset of overt motor symptoms up to 3 days before the onset of disease can be detected by conventional EAE scoring. This indicates that even subtle motor symptoms that arise during this stage^48^, which are presumably related to the formation of the first CNS lesions, can be detected by the ALMA toolbox. Our toolbox thus facilitates studies of the prodromal stage of the disease and its targeted modulation by therapeutic interventions.

Taken together we provide a user-friendly open-source toolbox that requires minimal time and resource commitments; is applicable to a wide range of neurological conditions affecting the brain and spinal cord; and provides an unbiased, robust, and comprehensive assessment of locomotion.

## Methods

### ALMA toolbox

This research aimed to provide a toolbox for the analysis of gait and footfall in mouse models of neurological disorders. This toolbox includes a graphical user interface (GUI) with functionalities for (i) automated kinematic parameter computation, (ii) automated footfall detection, (iii) data analysis of the computed kinematic parameters with random forest classification and principal component analysis, and (iv) visualization of gait kinematics. The ALMA toolbox is an open-source Python repository for the automatic processing of DLC coordinates, gait cycle detection, and kinematic parameter extraction. Following kinematic parameter extraction, the ALMA toolbox enables the use of machine learning algorithms, such as random forest classification and principal component analysis, to reduce dimensionality and identify the most relevant kinematic parameters in the scope of heath and disease. In addition to automated kinematic analysis, the toolbox GUI provides a code for the accurate detection of footfalls during fine motor tasks (e.g., traversing ladder rungs) for manual validation of each footfall. Details and the open-source toolbox can be found at https://github.com/sollan/alma

### Feature labeling and model training

To train the DLC model, we used approximately 450 image frames from different disease models (spinal cord injury, traumatic brain injury, and EAE) and timepoints. To predict hindlimb kinematic positions, we manually labeled six different body parts (toe, MTP joint, ankle, knee, hip, and iliac crest) in all ~450 image frames. The model was trained for up to 650,000 iterations using a deep residual network structure (ResNet-50), based on the pretrained model weights from DLC. To detect footfalls in the ladder rung test, we manually labeled all four paws on approximately 200 image frames for different mouse models (see above) and time points. The ResNet-50 model was trained for 400,000 iterations, based on the pretrained model weights from DLC. To train our models, we used a computer with 64 GB RAM, AMD Ryzen 9 3900X 12-Core Processor x 24, and GeForce RTX 2080 Ti 11 GB graphics card. Our trained models for kinematic and footfall analyses are publicly available.

### Animals

We used C57bl6 female mice of 2 to 4 months of age. The mice were kept in our animal facility under a regular day/night cycle (12 h/12 h). Animals had constant access to food and water. All animal experiments were carried out in accordance with the German animal welfare guidelines and previously authorized by the local regulatory committees (Regierung von Oberbayern).

### Animal Models and Surgeries

#### Spinal Cord Injury

Mice were anesthetized with MMF (medetomidin 0.5 mg/kg, Orion Pharma; midazolam 5.0 mg/kg, Ratiopharm; fentanyl 0.05 mg/kg, B. Braun). Once the mice presented no reflex reaction from paw pinching, their backs were shaved and a laminectomy was performed at T8 levels. The dura was exposed, and a dorsal hemisection was performed using irridectomy scissors^33,49^ (Bradley et al., 2019; Loy et al., 2018). After the hemisection, the wound was closed, and the skin was sutured. An antagonist mix was given (atipamezole 2.5 mg/kg, flumazenil 0.5 mg/kg, and naloxon 1.2 mg/kg), and mice were kept on a heating pad until completely awake. Mice received meloxicam (Metacam, 1.5 mg/ml oral suspension) at 12, 24, and 48 h following the injury.

#### EAE

Active EAE was induced in female mice by subcutaneously injection of 400 μg of purified recombinant MOG (N1-125) in complete Freund’s adjuvant (freshly made by adding 10 mg/mL *Mycobacterium* tuberculosis H37 Ra, Sigma-Aldrich). Then, 400 ng of pertussis toxin (Sigma-Aldrich) were administered intraperitoneally (i.p.) on days 0 and 2 after immunization^50^ (Locatteli et al., 2018). The mice were weighed daily and scored for neurological deficits according to the following EAE scores: 0, no clinical signs; 0.5, partial tail weakness; 1, tail paralysis; 1.5, gait instability or impaired righting ability; 2, hind limb paresis; 2.5, hind limb paresis with dragging of one foot; 3, total hind limb paralysis; 3.5, hind limb paralysis and forelimb paresis; 4, hind limb and fore limb paralysis; 5, death.

Stages of the disease were defined as follows: disease onset was defined as the first day with clinical symptoms in the EAE score, disease peak occurred 3 or 4 days after onset, and recovery occurred 10 or 11 days after onset).

#### Traumatic Brain Injury

Mice were anesthetized by intraperitoneal MMF injection. Once the mice presented no reflex reaction from paw pinching, they were put on a stereotaxic frame (Precision Systems & Instrumentation, LLC). A skin incision was made followed by the drilling of a window on the right skull hemisphere, which was positioned rostro-caudally, between the bregma and lambda and under the sagittal suture. Mice were then placed on the TBI-0310 impactor (Precision Systems & Instrumentation, LLC) to undergo traumatic brain injury (TBI). The tip of a 3-mm diameter steel rod was used to induce injury to the somatosensory cortex according to the following settings: 6 m/s, 150 ms dwell time, 0.5 mm depth^51^ (Littlejohn et al., 2020). Mice were removed from the impactor, the skull window was repositioned and sealed with Vetbond glue (3M Vetbond, 3M United States), and the skin was stitched. Mice were placed on a heating pad and injected with the antagonist mix before receiving a subcutaneous glucose injection (glucose 5% B. Braun Infusionslösung).

### Behavior Setup and Testing

#### Kinematics

Mice were recorded using a GoPro 8 camera at 120 frames per second while running on a treadmill (Harvard Apparatus; speed varying from 2 cm/s to 25 cm/s depending on the disease model; **Extended Fig. 1**). The distance between the treadmill and camera was 14.5 cm, and the camera was placed equidistant from the two ends of the treadmill. We chose this camera position to obtain comprehensive information on joint angles, spatial variability, and limb endpoint trajectories, which are missing from bottom recordings of gait analysis. Each mouse was recorded for at least 1 minute to sample enough step cycles. Every incomplete step cycle was automatically excluded by the toolbox. To extract meaningful datasets, a minimum number of 1,200 frames were captured, i.e., 10 s of recording. One prerequisite to accurate data extraction was the blind selection of frames with representative step cycles (no pause in animal locomotion, no grooming, no back turns etc.). When the animal’s limbs are completely dragging, our algorithm will only compute parameters that are independent to the step cycles. All preprocessing and processing steps undertaken by the ALMA toolbox can be found in **Extended Fig. 1**.

#### Ladder Rung

Mice were recorded using a GoPro 8 camera at 120 frames per second while making four consecutive runs on our custom-made ladders. In this test, the animals had to cross 1-m horizontal ladders, and footfalls were recorded. We evaluated the rhythmic locomotion in the regular walking task on a ladder with evenly spaced rungs and the animal’s fine coordination paw placement ability using irregularly spaced rungs (irregular walking task). The camera was placed 17.5 cm distant from the ladder equidistant from the two ends. Animals were habituated on a ladder with regularly spaced rungs before any experiment was performed (2/3 habituations each for a total max. time of 3 minutes). We used DLC to perform markerless labeling of the hind paws and then applied ALMA to determine footfall characteristics with a peak detection algorithm. In the toolbox, we included three preprocessing algorithms that accommodate a range of different recording conditions and can be chosen by the experimenter. Here we chose to use a preprocessing procedure based on the deviation algorithm that shows a great correlation with manual counting. All data preprocessing and analysis steps undertaken by the ALMA toolbox can be found in **Extended Fig. 2**.

#### Basso Mouse Scale

All mice with SCI were evaluated pre-operatively and at 7 and 21dpi. The scores regarding the locomotor ability of mice were given according to the original paper^27^ by fully trained observers.

### Statistical Analysis

All results are given as mean ± standard error of the mean (SEM) in the figures. In the expanded tables, data are given as mean ± standard deviation (SD). Statistical analysis and the construction of the graphs for data illustration were carried out on GraphPad Prism 8.4.3 for Windows (GraphPad Software). All datasets were tested for normality using the Shapiro–Wilk test. Parametric data from the SCI and TBI experiments were analyzed using repeated measures ANOVA and Tukey’s post-hoc test. Nonparametric datasets from the SCI and TBI experiments were analyzed using the Friedman test followed by the Dunn post-hoc test. Datasets from the EAE experiments were analyzed with the Kruskal–Wallis test followed by the Dunn post-hoc test, as they were distributed non-normally. To determine correlation, we used the Pearson’s correlation coefficient. In order to classify animals, we used random forest classification, based on individual step cycles. Feature importance in the Random Forest Classification was determined by Gini impurity-based feature importance which ranged between 0 and 1. Statistical significance levels are indicated as follows: *p < 0.05; **p < 0.01; ***p < 0.001.

## Acknowledgments

The authors would like to thank Luca Fabbio for animal handling as well as Dana Matzek and Bianca Stahr for animal husbandry. Work in F.M.B.’s lab is supported by grants from the Deutsche Forschungsgemeinschaft (DFG, SFB 870 Project ID 118803580, TRR274 Project ID 408885537), by the Munich Center for Neurosciences (MCN) and the Institute for Research on Paraplegia (IRP). F.M.B. is also supported by the Munich Center for Systems Neurology (DFG, SyNergy; EXC 2145 / ID 390857198). VVS is supported by a post-doctoral fellowship from the Humboldt foundation. The authors declare no competing financial interests.

## Author contributions

FB, AA, and SZ designed the experiments. AA, CRV, and MC performed all surgical procedures. AA, SZ and VV collected and analyzed data. FB, AA, SZ, and MK wrote the manuscript. All authors approved the final version of the paper.

## Code availability

We have provided the code for the ALMA toolbox at https://github.com/sollan/alma

---

**Extended Figure 1:**
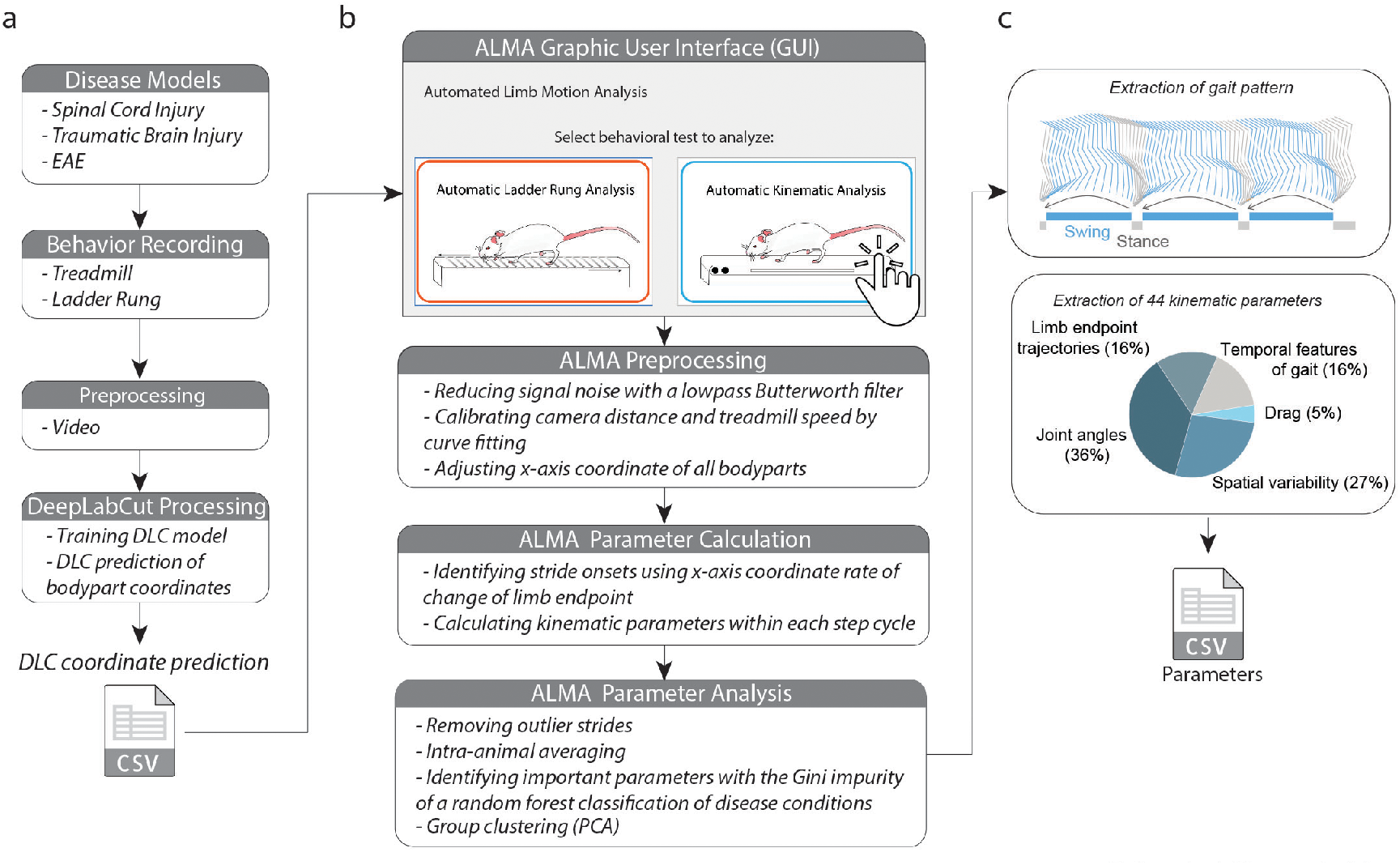
Detailed flowchart of the ALMA computations used to obtain kinematic parameters. **a**- Flowchart of the experiment, including selection of injury paradigm, behavior recording, video preprocessing, DeepLabCut (DLC) markerless labeling, model training, and coordinate export. **b-** Workflow of the ALMA toolbox, with a user-friendly graphical user interface (GUI) for choosing kinematic analysis, and the computational steps used to identify important parameters and perform group clustering. **c-** ALMA allows the user to extract gait patterns and analyze the 44 parameters.

**Extended Figure 2:**
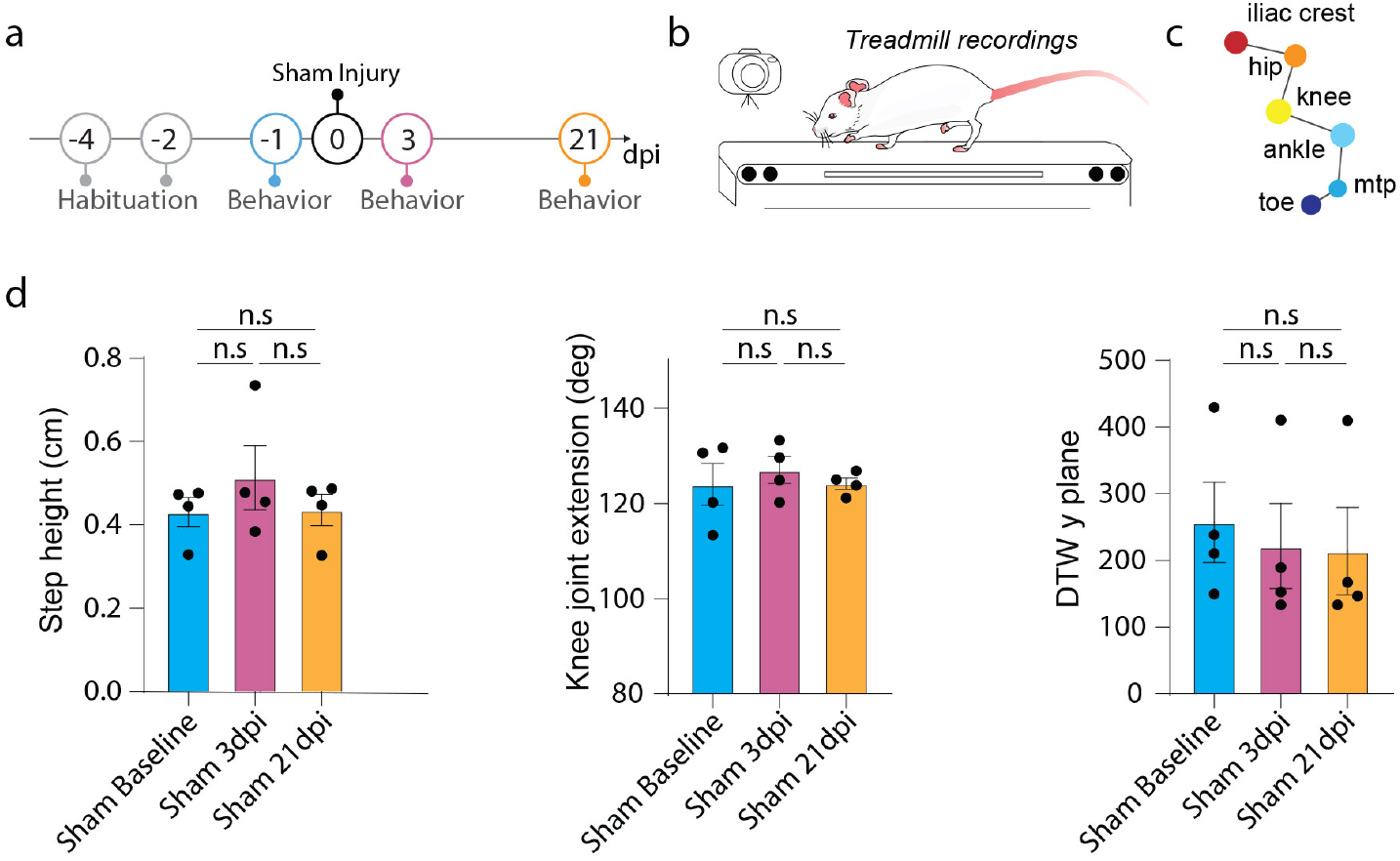
ALMA analysis does not detect any changes of gait in sham injured mice tested on the treadmill. **a-** Timeline of the sham experiment. **b-** Schematic of the treadmill system used to record the behavior of mice. **c-** Schematic of DeepLabCut (DLC) markerless joint labeling. Six joints were labeled: iliac crest, hip, knee, ankle, metatarsophalangeal joint (MTP), and toe. **d-** Quantitative evaluation of the same parameters as in Figure 2, such as step height, knee joint extension, or dynamic time warping (DTW), at baseline, 3 and 21 days post-sham surgery. Repeated one-way ANOVA was used to analyze knee joint extension (p = 0.440; n = 4), step height (p=0.458; n=4) and DTW y plane (p=850; n = 4). In all panels, data are presented as mean± SEM; *p < 0.05; **p < 0.01; ***p < 0.001. dpi: days post-injury.

**Extended Figure 3:**
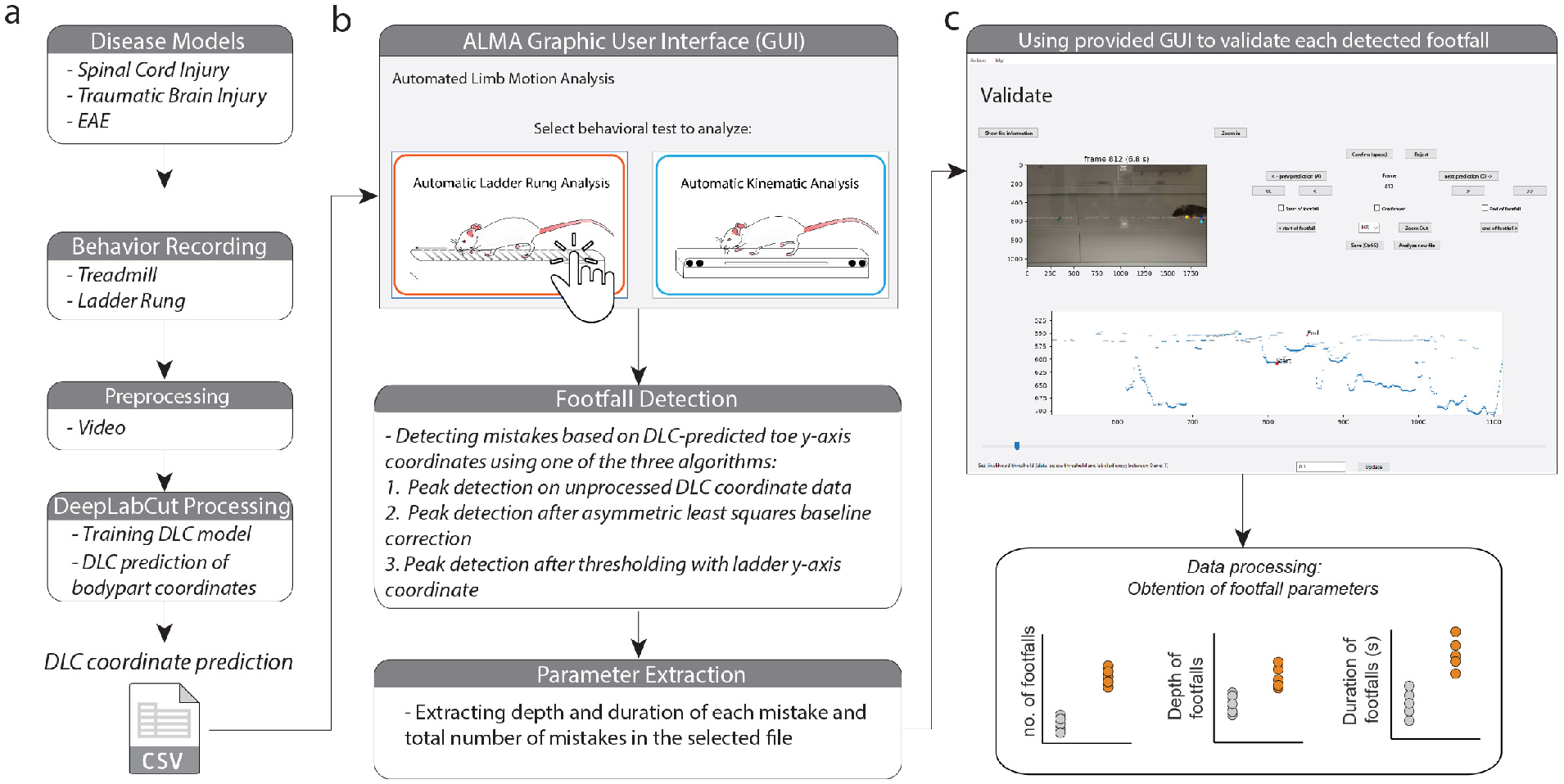
Detailed flowchart of ALMA computations used to obtain footfall parameters. **a**- Flowchart of the experiment, including choice of injury paradigm, behavioral recording, video preprocessing, DeepLabCut (DLC) markerless labeling, model training, and coordinate export. **b-** Workflow of the ALMA toolbox, with a user-friendly graphical user interface (GUI) for selecting ladder rung analysis, and computational steps to identify the number, depth, and duration of footfalls. **c-** Each detected footfall can be visualized and manually validated or excluded before final data are generated by ALMA.

**Extended Table 1:**
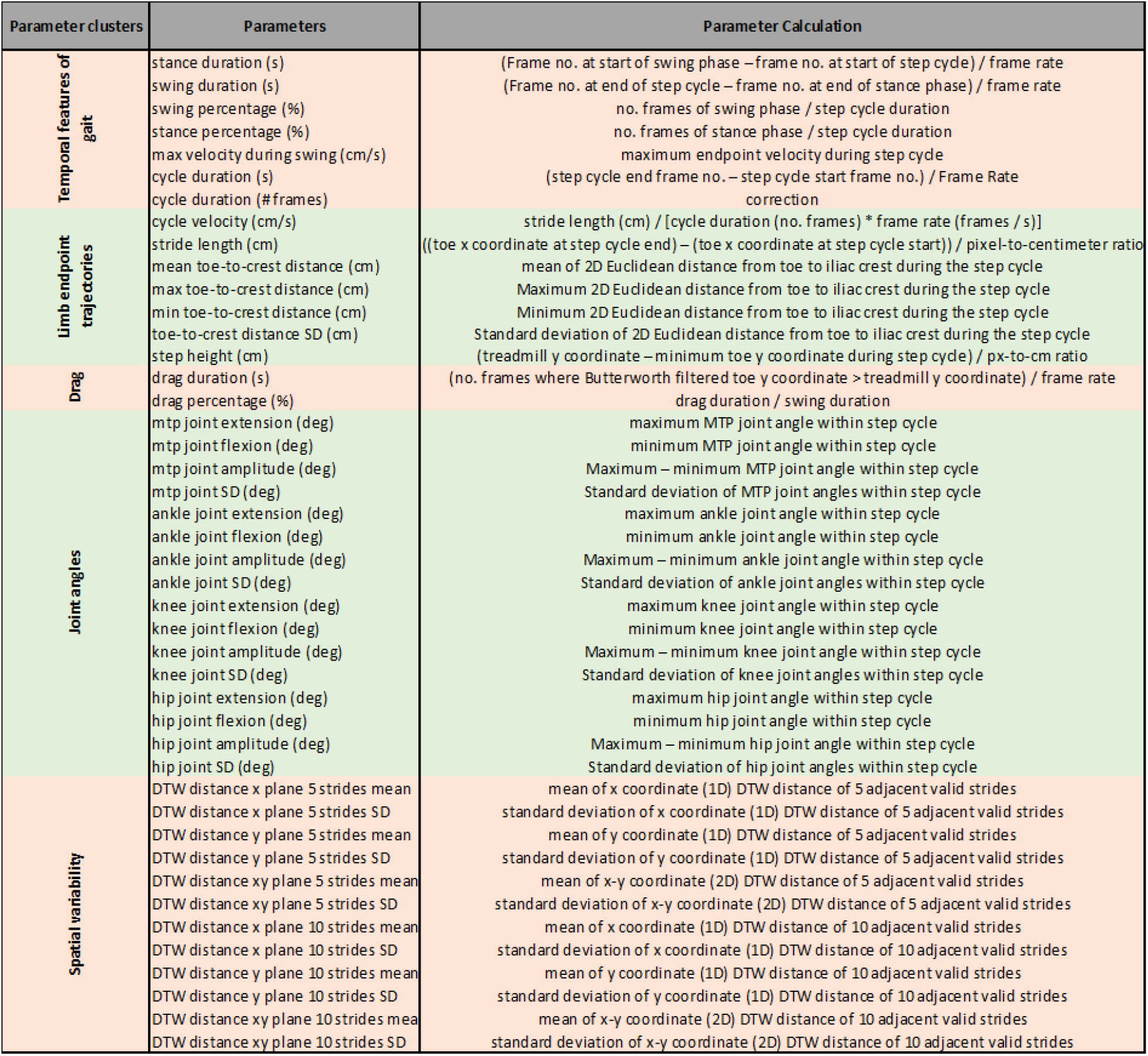
Description and mathematical formulae for parameters calculated by the ALMA toolbox.

**Extended Table 2:** Quantitative measurement of 44 spinal cord injury kinematic parameters obtained by ALMA at baseline, 3 dpi, and 21 dpi. Data are presented as mean ± SD.

**Extended Table 1: Description and mathematical formulae for parameters calculated by the ALMA toolbox.**
| Parameter clusters | Parameters | Baseline |  | 1DPI |  | 10DPI |  |
| --- | --- | --- | --- | --- | --- | --- | --- |
|  |  | Mean | SD | Mean | SD | Mean | SD |
| Temporal features of gait | stance duration (s) | 0.272501 | 0.02778 | 0.324468 | 0.073975 | 0.278786 | 0.070118 |
|  | swing duration (s) | 32.66743 | 3.330316 | 38.89727 | 8.868065 | 33.42085 | 8.405765 |
|  | swing percentage (%) | 20.01785 | 2.378841 | 13.24887 | 6.097586 | 17.69423 | 5.598283 |
|  | stance percentage (%) | 5.030653 | 0.438425 | 3.736301 | 1.115117 | 4.370403 | 0.794919 |
|  | max velocity during swing (cm/s) | 0.128701 | 0.022238 | 0.154144 | 0.038008 | 0.131397 | 0.029552 |
|  | cycle duration (s) | 0.1438 | 0.01839 | 0.170325 | 0.04709 | 0.147389 | 0.04209 |
|  | cycle duration (no. frames) | 0.531433 | 0.05752 | 0.521877 | 0.040124 | 0.519481 | 0.034092 |
| Limb endpoint trajectories | cycle velocity (cm/s) | 0.468567 | 0.05752 | 0.478123 | 0.040124 | 0.480519 | 0.034092 |
|  | stride length (cm) | 2.479889 | 0.449962 | 2.395532 | 0.286082 | 2.627573 | 0.375719 |
|  | mean toe-to-crest distance (cm) | 3.053605 | 0.345247 | 2.819122 | 0.346363 | 3.044387 | 0.365592 |
|  | max toe-to-crest distance (cm) | 2.006263 | 0.492077 | 2.083746 | 0.247717 | 2.274772 | 0.372092 |
|  | min toe-to-crest distance (cm) | 0.316346 | 0.065982 | 0.226115 | 0.059471 | 0.239906 | 0.094296 |
|  | toe-to-crest distance SD (cm) | 0.418506 | 0.082368 | 0.129611 | 0.071069 | 0.357871 | 0.253824 |
|  | step height (cm) | 53.82473 | 4.310894 | 41.64993 | 16.02726 | 48.72549 | 9.950884 |
| Drag | drag duration (s) | 172.4974 | 4.808273 | 165.2026 | 9.880856 | 169.1852 | 4.10966 |
|  | drag percentage (%) | 110.9118 | 25.12176 | 111.1831 | 8.887672 | 120.9324 | 13.46444 |
| Joint angles | mtp joint extension (deg) | 61.58558 | 21.94498 | 54.01952 | 8.439814 | 48.25288 | 13.17755 |
|  | mtp joint flexion (deg) | 21.04034 | 9.049714 | 17.2016 | 2.713314 | 15.7355 | 4.239146 |
|  | mtp joint amplitude (deg) | 115.2692 | 5.197123 | 127.5739 | 19.01481 | 115.0247 | 10.9949 |
|  | mtp joint SD (deg) | 66.17388 | 14.55343 | 72.41404 | 20.717 | 68.1328 | 12.41157 |
|  | ankle joint extension (deg) | 49.09534 | 16.15845 | 55.15981 | 12.69716 | 46.89194 | 9.688858 |
|  | ankle joint flexion (deg) | 16.1417 | 4.982632 | 18.40307 | 4.211443 | 15.79921 | 3.228546 |
|  | ankle joint amplitude (deg) | 126.2849 | 6.819535 | 103.8706 | 13.36608 | 126.2695 | 6.818621 |
|  | ankle joint SD (deg) | 84.43803 | 15.08882 | 76.30352 | 13.38735 | 95.49433 | 8.506513 |
|  | knee joint extension (deg) | 41.84686 | 14.68188 | 27.56712 | 4.866101 | 30.77513 | 11.23552 |
|  | knee joint flexion (deg) | 14.05208 | 4.855436 | 8.731922 | 1.816848 | 9.909117 | 3.596147 |
|  | knee joint amplitude (deg) | 101.0559 | 15.50737 | 98.99893 | 10.52836 | 108.8717 | 16.53888 |
|  | knee joint SD (deg) | 59.93486 | 6.361746 | 65.05851 | 11.32375 | 73.21607 | 6.771436 |
|  | hip joint extension (deg) | 41.12103 | 15.67308 | 33.94042 | 10.02455 | 35.65567 | 12.97713 |
|  | hip joint flexion (deg) | 14.0558 | 5.617646 | 11.42244 | 3.801759 | 12.06421 | 4.75993 |
|  | hip joint amplitude (deg) | 0.002212 | 0.001304 | 0.008046 | 0.008611 | 0.003418 | 0.003746 |
|  | hip joint SD (deg) | 0.004198 | 0.002956 | 0.044267 | 0.049751 | 0.014499 | 0.019797 |
| Spatial variability | DTW distance x plane 5 strides mean | 846.4958 | 459.622 | 881.9535 | 240.2091 | 825.1326 | 359.3023 |
|  | DTW distance x plane 5 strides SD | 672.7041 | 528.8476 | 672.214 | 280.0555 | 621.6206 | 364.4608 |
|  | DTW distance y plane 5 strides mean | 390.5996 | 356.6238 | 187.8594 | 208.1919 | 336.4633 | 414.1873 |
|  | DTW distance y plane 5 strides SD | 366.6243 | 419.538 | 165.2883 | 231.8537 | 281.3317 | 412.1286 |
|  | DTW distance xy plane 5 strides mean | 1114.693 | 641.9847 | 1011.349 | 295.3583 | 1064.549 | 581.26 |
|  | DTW distance xy plane 5 strides SD | 842.2055 | 728.3903 | 745.0359 | 348.713 | 744.0106 | 551.4335 |
|  | DTW distance x plane 10 strides mean | 850.8711 | 467.0968 | 847.5654 | 241.5939 | 852.2332 | 403.8618 |
|  | DTW distance x plane 10 strides SD | 932.5275 | 836.09 | 836.4553 | 423.1833 | 862.8633 | 607.68 |
|  | DTW distance y plane 10 strides mean | 393.4395 | 361.2572 | 189.993 | 211.1147 | 345.7809 | 439.7726 |
|  | DTW distance y plane 10 strides SD | 540.5131 | 687.5086 | 243.453 | 377.3961 | 414.5704 | 667.7815 |
|  | DTW distance xy plane 10 strides mean | 1120.609 | 654.5719 | 979.3329 | 302.8977 | 1097.534 | 629.7265 |
|  | DTW distance xy plane 10 strides SD | 1180.169 | 1176.488 | 950.7398 | 559.1539 | 1044.252 | 898.6512 |

**Extended Table 3:** Quantitative measurement of 44 traumatic brain injury kinematic parameters obtained by ALMA at baseline, 1 dpi, and 10 dpi. Data are presented as mean ± SD.

| Parameter clusters | Parameters | Baseline |  | 1DPI |  | 10DPI |  |
| --- | --- | --- | --- | --- | --- | --- | --- |
|  |  | Mean | SD | Mean | SD | Mean | SD |
| Temporal features of gait | stance duration (s) | 0.121824 | 0.015986 | 0.118299 | 0.013777 | 0.120694 | 0.010487 |
|  | swing duration (s) | 0.114017 | 0.009108 | 0.132097 | 0.019376 | 0.119472 | 0.013917 |
|  | swing percentage (%) | 0.488084 | 0.036879 | 0.520157 | 0.040942 | 0.494563 | 0.028555 |
|  | stance percentage (%) | 0.511916 | 0.036879 | 0.479843 | 0.040942 | 0.505437 | 0.028555 |
|  | max velocity during swing (cm/s) | 60.76525 | 4.108749 | 56.56078 | 4.905492 | 59.04529 | 5.090353 |
|  | cycle duration (s) | 0.235841 | 0.018705 | 0.250396 | 0.020603 | 0.240166 | 0.021154 |
|  | cycle duration (no. frames) | 28.27267 | 2.242367 | 30.0175 | 2.469888 | 28.79111 | 2.535994 |
| Limb endpoint trajectories | cycle velocity (cm/s) | 24.03619 | 1.860084 | 21.56867 | 2.675134 | 23.06046 | 2.2548 |
|  | stride length (cm) | 5.467224 | 0.185781 | 5.049387 | 0.356628 | 5.303987 | 0.263472 |
|  | mean toe-to-crest distance (cm) | 2.697055 | 0.07725 | 2.629069 | 0.187952 | 2.749006 | 0.154402 |
|  | max toe-to-crest distance (cm) | 3.040595 | 0.085559 | 3.046766 | 0.208821 | 3.128813 | 0.175387 |
|  | min toe-to-crest distance (cm) | 2.308645 | 0.08491 | 2.265401 | 0.166957 | 2.341604 | 0.170356 |
|  | toe-to-crest distance SD (cm) | 0.226423 | 0.024947 | 0.233403 | 0.029264 | 0.243264 | 0.027067 |
|  | step height (cm) | 0.470995 | 0.033842 | 0.364227 | 0.064624 | 0.454162 | 0.045857 |
| Drag | drag duration (s) | 0.002077 | 0.003029 | 0.005172 | 0.004947 | 0.002338 | 0.001843 |
|  | drag percentage (%) | 0.003694 | 0.005337 | 0.012383 | 0.00921 | 0.004401 | 0.003797 |
| Joint angles | mtp joint extension (deg) | 175.5036 | 1.753417 | 174.4127 | 2.896869 | 173.6816 | 4.232018 |
|  | mtp joint flexion (deg) | 123.8531 | 5.63525 | 121.8831 | 8.1265 | 119.2426 | 2.745881 |
|  | mtp joint amplitude (deg) | 51.65052 | 5.020858 | 52.52962 | 6.924255 | 54.43905 | 5.164657 |
|  | mtp joint SD (deg) | 17.18208 | 1.634765 | 17.42373 | 2.517821 | 18.12524 | 1.821725 |
|  | ankle joint extension (deg) | 113.137 | 3.532662 | 110.5983 | 4.017837 | 116.4725 | 3.75105 |
|  | ankle joint flexion (deg) | 72.58927 | 6.516049 | 70.93998 | 4.489512 | 71.86161 | 3.429244 |
|  | ankle joint amplitude (deg) | 40.54775 | 6.41054 | 39.65832 | 5.788168 | 44.61088 | 1.698437 |
|  | ankle joint SD (deg) | 13.77948 | 2.039465 | 13.10895 | 2.07707 | 15.14024 | 0.585989 |
|  | knee joint extension (deg) | 126.6993 | 3.974741 | 125.9935 | 4.241151 | 125.8478 | 6.535263 |
|  | knee joint flexion (deg) | 97.21903 | 6.4127 | 93.84855 | 2.810883 | 93.93848 | 5.141963 |
|  | knee joint amplitude (deg) | 29.48023 | 3.394054 | 32.14492 | 5.872531 | 31.90928 | 3.259213 |
|  | knee joint SD (deg) | 10.19764 | 1.280341 | 10.94247 | 1.958866 | 10.78471 | 1.174599 |
|  | hip joint extension (deg) | 103.7711 | 5.299289 | 102.7507 | 5.37069 | 103.9755 | 5.317694 |
|  | hip joint flexion (deg) | 66.78887 | 4.757523 | 68.67242 | 2.336178 | 68.20187 | 4.316921 |
|  | hip joint amplitude (deg) | 36.98223 | 4.181746 | 34.07829 | 5.372466 | 35.77363 | 5.799337 |
|  | hip joint SD (deg) | 12.54367 | 1.501702 | 11.5638 | 2.17545 | 12.27671 | 2.055408 |
| Spatial variability | DTW distance x plane 5 strides mean | 540.3867 | 212.6678 | 917.2769 | 398.675 | 534.6435 | 104.9336 |
|  | DTW distance x plane 5 strides SD | 327.3784 | 158.1207 | 733.6116 | 448.5399 | 324.158 | 116.3877 |
|  | DTW distance y plane 5 strides mean | 166.5777 | 69.33384 | 437.2623 | 299.5536 | 180.2077 | 101.5493 |
|  | DTW distance y plane 5 strides SD | 121.3037 | 67.77559 | 431.9553 | 348.5609 | 119.0592 | 81.1103 |
|  | DTW distance xy plane 5 strides mean | 678.5163 | 252.6987 | 1203.494 | 534.4544 | 700.3912 | 178.0202 |
|  | DTW distance xy plane 5 strides SD | 373.6489 | 178.48 | 941.5367 | 599.772 | 378.0759 | 135.8761 |
|  | DTW distance x plane 10 strides mean | 506.2561 | 178.4697 | 864.9111 | 331.0459 | 507.8059 | 99.62951 |
|  | DTW distance x plane 10 strides SD | 373.0266 | 190.7426 | 937.0467 | 579.7646 | 386.655 | 154.4982 |
|  | DTW distance y plane 10 strides mean | 160.743 | 69.26296 | 421.6389 | 268.3219 | 179.0769 | 98.20865 |
|  | DTW distance y plane 10 strides SD | 156.472 | 104.2623 | 618.8024 | 497.2211 | 144.4497 | 102.226 |
|  | DTW distance xy plane 10 strides mean | 639.7849 | 222.8888 | 1146.656 | 459.0672 | 674.2344 | 179.8192 |
|  | DTW distance xy plane 10 strides SD | 435.1609 | 228.9883 | 1244.121 | 801.8841 | 455.6019 | 177.9623 |

**Extended Table 4:**
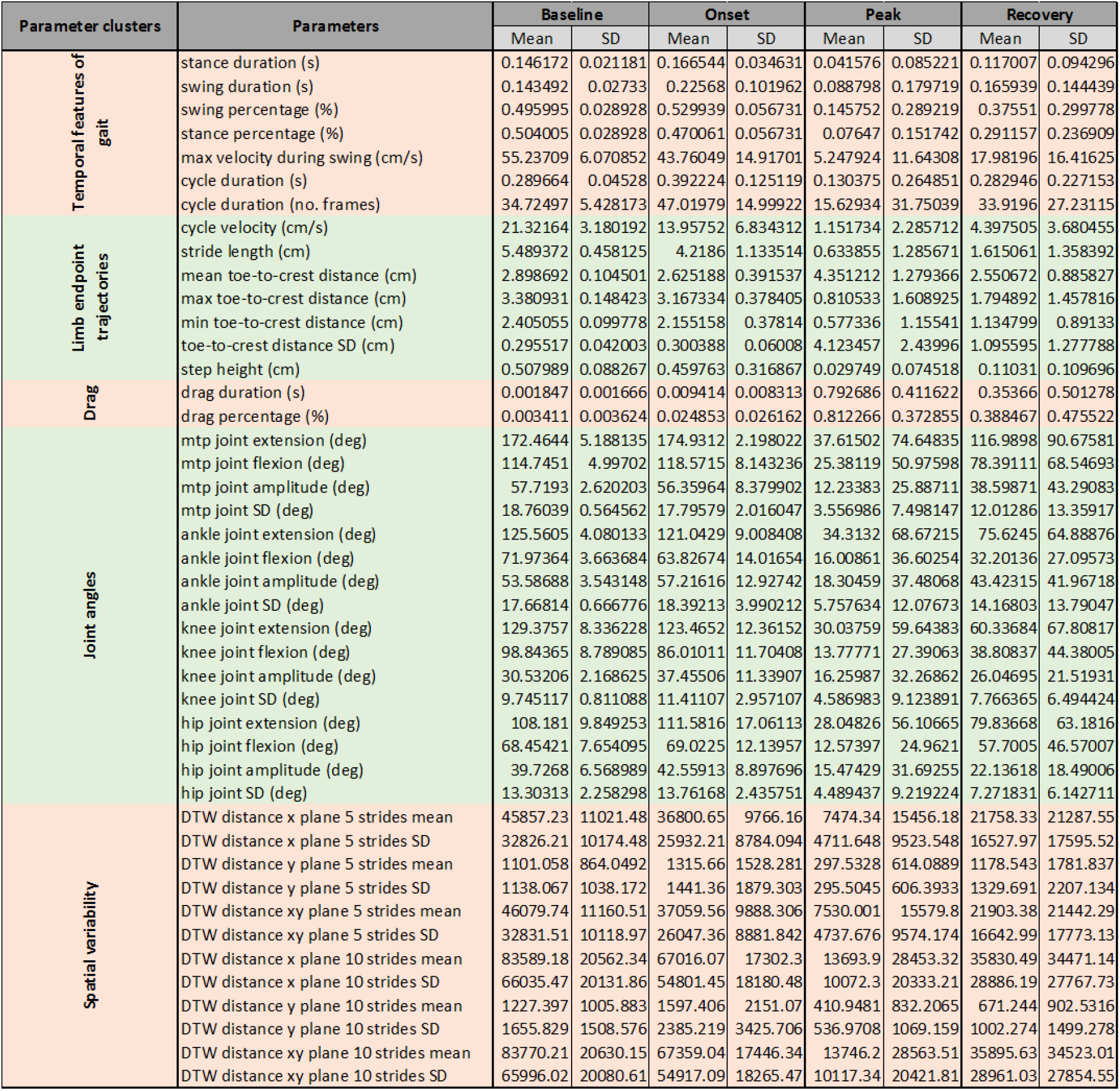
Quantitative measurement of 44 EAE kinematic parameters obtained by ALMA at baseline, onset, peak, and disease recovery. Data are presented as mean ± SD.

## Notes

https://github.com/sollan/alma/

### Competing Interest Statement

The authors have declared no competing interest.

